# Proteolytic Processing of the Unique F_1_-ATPase Subunit α in Procyclic *Trypanosoma brucei*

**DOI:** 10.1101/2025.10.24.684300

**Authors:** Jain Minal, Panicucci Brian, Šubrtová Karolína, Schnaufer Achim, Zíková Alena

**Affiliations:** Biology Centre, Institute of Parasitology, České Budějovice, Czech Republic; Faculty of Science, University of South Bohemia, České Budějovice, Czech Republic; Institute of Immunology and Infection Research, University of Edinburgh, Edinburgh, United Kingdom

## Abstract

The catalytic F_1_ domain of the mitochondrial ATP synthase is conserved across eukaryotes, with the only known exceptions found within euglenozoans. One distinctive trait identified in *Trypanosoma brucei* is the proteolysis of subunit α that excises an internal octapeptide, resulting in functional N- and C-terminal polypeptides. The significance of this cleavage remained unclear. Here, we determined that the yeast subunit α expressed in *T. brucei* was not proteolytically processed, despite significant structural similarities. The proteolytic recognition sequence was identified to be largely contained within the octapeptide and replacing it with a flexible linker rendered the protein resistant to cleavage. This uncleaved subunit α restored growth after RNAi depletion of endogenous subunit α by incorporating into F_1_F_o_-ATP synthase complexes capable of both ATP synthesis and hydrolysis. A FRET-based assay revealed that peptides consisting of either the first or second octapeptide cleavage site experienced significantly more proteolysis when incubated with cytosolic lysates compared to mitochondrial extracts. Furthermore, expressing just the mature C-terminal polypeptide resulted in mitochondrial localization, suggesting it contains an internal targeting signal. Together, these results indicate that proteolysis of subunit α occurs in the cytosol prior to mitochondrial import, highlighting a unique processing step in *T. brucei* ATP synthase assembly.

## INTRODUCTION

Generally, aerobic eukaryotes produce most of their cellular energy in the form of ATP through oxidative phosphorylation. This process relies on reduced cofactors produced during the catabolism of energy sources to provide electrons to a series of protein complexes residing within the mitochondrial cristae. As electrons flow through the electron transport chain, the energy released from these reduction-oxidation reactions is used to pump protons across the mitochondrial inner membrane. The potential energy from this proton motive force is then utilized by the F_1_F_O_-ATP synthase to generate kinetic energy as protons allowed to flow down their gradient into the mitochondrial matrix propel the rotation of this elegant nanomotor. This proton pore is formed between the oligomeric subunit *c*-ring and subunit *a*, both of which are localized to the membrane embedded F_O_ domain. As the protons move through these half-channels, static charges drive the rotation of the c-ring [1]. This in turn rotates the firmly attached central rotor stalk subunit γ of the catalytic F_1_-ATPase domain, a multiprotein structure that protrudes into the mitochondrial matrix. Each 120° rotation by the asymmetrical subunit γ causes conformational changes within the catalytic nucleotide binding sites located at the interface of three heterodimers of alternating α and β subunits [2], resulting in three catalytic states: open (low affinity, releases ATP), loose (binds ADP + P_i_) and tight (ATP formation) [3]. Thus, after one complete 360° rotation of subunit γ, three ATP molecules are synthesized from ADP and inorganic phosphate. Importantly, the subunit α and β hexameric headgroup is fixed in place through the binding of one subunit α with the oligomycin sensitivity conferral protein (OSCP) of the stator stalk, a peripheral arm typically anchored to subunit *a* in the F_O_ domain [4].

The composition and structure of the mitochondrial F_1_F_o_-ATP synthase can vary across eukaryotic lineages. For example, the number of subunit *c* proteins in the *c*-ring ranges from eight in bovine [5] to ten in yeast [6], altering the proton-to-ATP ratio such that enzymes with fewer *c* subunits are more efficient at synthesizing ATP [7]. Although all mitochondrial F_1_F_o_-ATP synthases form dimers through contacts between the F_o_ moieties, the dimerization interface differs between species, in part due to lineage-specific subunits [8–11]. Cryogenic electron microscopy and tomography studies have identified at least four different types of dimers [12], distinguished by the angle between the central stalks and the orientation of the paired monomers respective to each other [13–21].These dimers self-assemble into rows that bend the membrane, helping to form the mitochondrial cristae [18]. These rows vary in their organization and length, resulting in either straight rows [18,22], uneven rows [23], long twisted rows [13] or short ribbons [21]. Thus, the structural diversity of ATP synthase dimers and rows contributes the distinct cristae morphologies observed across eukaryotes [24–26].

While there are important differences in the F_o_ domain across eukaryotic lineages, the F_1_-ATPase domain was long considered to be highly conserved in composition and structure. However, some remarkable features were identified in the euglenozoan F_1_-ATPase [27,28]. Euglenozoa are a large group of flagellated protists that include free-living species along with several important parasites, including *Trypanosoma brucei*, the etiological agent of human and animal African trypanosomiases in sub-Saharan Africa. Due to the ease of cultivation and genetic manipulation, *T. brucei* is a model organism with an extensive informatics online platform [29–31]. Moreover, the parasite provides an excellent tool to study mitochondrial biogenesis as the organelle undergoes structural remodeling and metabolic rewiring during the parasite development in mammalian and insect hosts. In the insect vector, the procyclic form (PF) has a single reticulated mitochondrion with abundant discoidal cristae containing a canonical electron transport chain (ETC) and an F_1_F_o_-ATP synthase that synthesizes ATP [32–35]. However, when the extracellular parasite resides in the glucose abundant mammalian bloodstream, it lacks ETC complexes required for oxidative phosphorylation and relies on glycolysis for ATP production. In the long slender bloodstream form (BF), the F_1_F_o_-ATP synthase continuously hydrolyzes ATP, which causes it to rotate in reverse and pump protons from the mitochondrial matrix into the intermembrane space [36]. This unique function is essential for maintaining the mitochondrial membrane potential in a mitochondrion that has been remodeled into a tubular structure characterized by limited cristae with reduced volume [37].

Independent of the life cycle stage, the *T. brucei* F_1_-ATPase is elaborated with three copies of the euglenozoan specific subunit p18. This pentatricopeptide repeat protein [38] associates with the external surface of subunit α, resulting in a non-canonical F_1_-ATPase head that resembles a triangular pyramid [21]. However, the function of p18 remains unknown as the enzyme appears to have a conventional catalytic mechanism for ATP production due to the highly conserved *T. brucei* F_1_-ATPase α and β subunits [27]. Both of these subunits are comprised of three domains, which are shown schematically for subunit α in Figure 1A. The N-terminal domain consists of a six-stranded β barrel that forms the crown of the F_1_-ATPase. Sitting atop of the peripheral stalk is subunit OSCP, which binds the crown of subunit α and stabilizes the F_1_-ATPase moiety. The central domain of subunit α and β is the nucleotide-binding domain, with the defined Walker A and B motifs. Finally, the C-terminal domain of these subunits is comprised of several α-helices.

**Figure 1.**
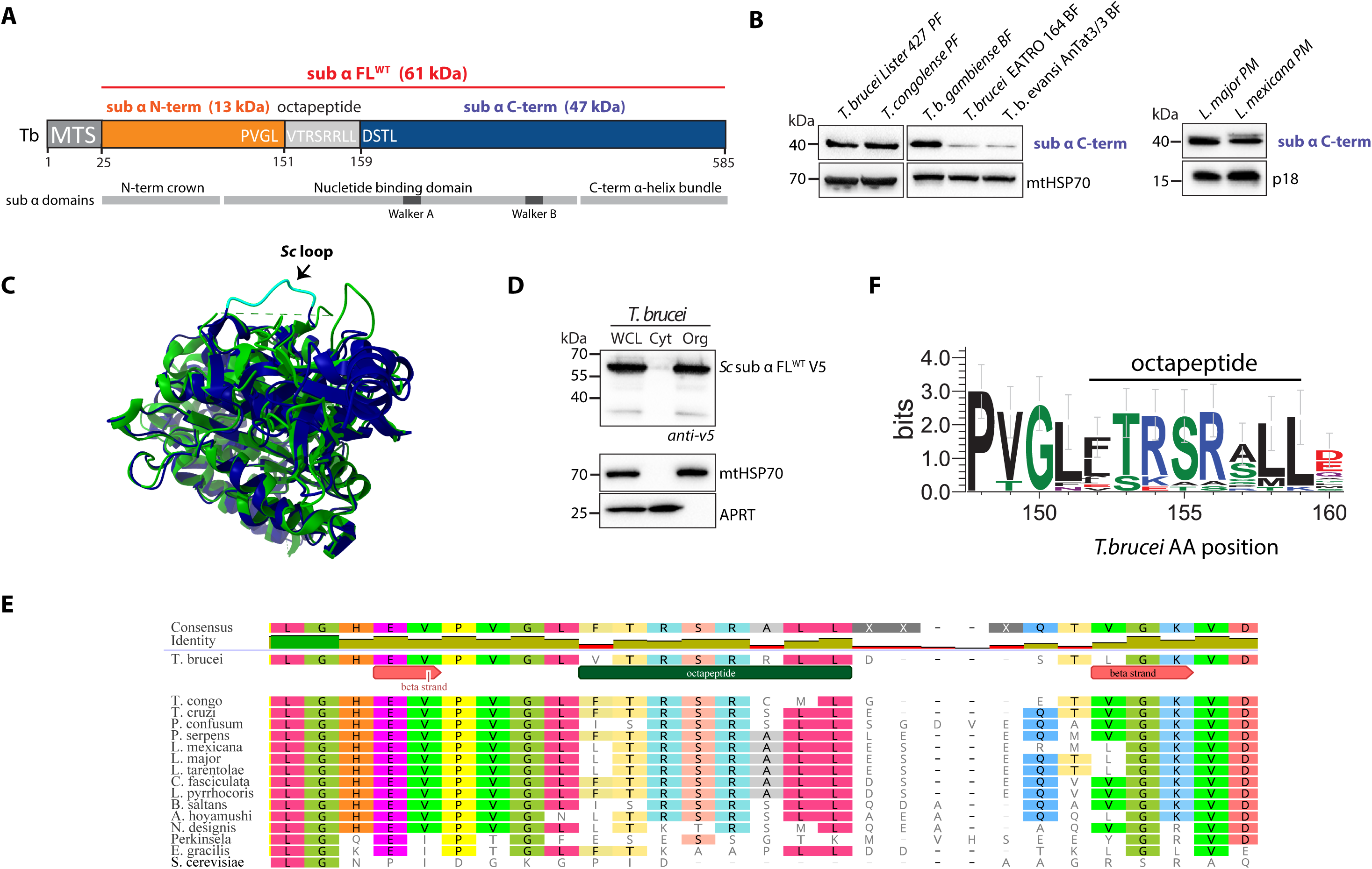
F_1_-ATPase subunit α is cleaved at specific sites across euglenozoan species. **A.** A schematic illustrating the polypeptides of the mature *T. brucei* sub α FL^WT^ including the mitochondrial targeting sequence (MTS) and the amino acid sequence of the excised octapeptide and adjacent residues. The numerical position of the amino acids immediately to the C-terminus of the cleaved peptides are indicated. Approximate locations of important domains identified in opisthokonts are depicted in grey bars. **B.** Immunoblot of mitochondrial lysates from several model Trypanosomatids probed with an antibody recognizing subunit α C-term. The relevant sizes of the protein marker are indicated on the left. PF-procyclic form, BF–bloodstream form, PM-promastigote. **C.** Superimposition of subunit α from *T. brucei* (6F5D, green) and *S. cerevisiae* (6CP6, blue). The root mean squared deviation (RMSD) between 368 pruned atom pairs is 0.907 angstroms. The yeast subunit α structure depicted in cyan corresponds to the excised *T. brucei* octapeptide. **D.** Western blot analysis of whole cell lysate (WCL), cytosol (Cyt) and digitonin-extracted enriched organellar fractions (Org) from *T. brucei* expressing yeast subunit α tagged with a V5 epitope. Specific antibodies recognizing either the *T. brucei* cytosolic APRT or mitochondrial (mt) HSP70 were used to detect the respective cellular compartments. The relevant sizes of the protein marker are indicated on the left. **E.** The Clustal Omega alignment is focused on the region around the *T. brucei* octapeptide (green box) that is proteolytically removed. Defined structural features determined from the *T. brucei* F_1_-ATPase X-ray crystallography studies [27] are depicted with red directional boxes. The gene sequences from the various euglenozoan species (*Trypanosoma congolense, Trypanosoma cruzi, Paratrypanosoma confusum, Phytomonas serpens, Leishmania mexicana, Leishmania major, Leishmania tarentolae, Crithidia fasciculata, Leptomonas pyrrhocoris, Bodo saltans, Azumiobodo hoyamushi, Neobodo designis, Perkinsela, Euglena gracisili*) and *Saccharomyces cerevisiae* are listed in descending order according to their increasing phylogenetic distance from *T. brucei*, which is based largely on 18S rRNA [74]. **F.** Sequence weblogo of the region around the *T. brucei* octapeptide based on the alignment of subunits α in **C**. The identical sequences from *T.b. gambiense* and *T.b. evansi* were removed from the analysis. Weblogo coloring: polar amino acids (AAs, green), neutral AAs (purple), basic AAs (blue), acidic AAs (red), hydrophobic AAs (black).

Another unique attribute identified in the euglenozoan F_1_-ATPase is the proteolytic processing of subunit α that results in mature N-terminal and C-terminal peptides [11,39–42]. The complexity of this proteolysis was later revealed when the F_1_-ATPase synthase was purified from *T. brucei* and the peptides were analyzed by Edman sequencing [43]. This determined that the *T. brucei* subunit α (TriTrypDB Gene IDs Tb927.7.7420 and Tb927.7.7430) is proteolytically cleaved three times (Figure 1A). The first cleavage occurs after amino acid 24, which removes the N-terminal mitochondrial targeting sequence (MTS). The other two proteolytic events occur after leucine residues 151 and 159. This results in the expulsion of an octapeptide, generating a mature N-terminal subunit α with a predicted molecular weight of 13 kDa and a C-terminal subunit of 47 kDa. In this study we investigate the biological significance of this cleavage in the cultured insect stage of *T. brucei*.

## RESULTS AND DISCUSSION

### The proteolytical processing of euglenozoan F_1_-ATPase subunit α is sequence specific

Subunit α proteolysis has been experimentally demonstrated only in the following euglenozoa: *Crithidia fasciculata* [39], *Phytomonas serpens* [40], *T. brucei brucei* [41], *Leishmania tarentolae* [42] and *Euglena gracilis* [11]. Therefore, to generate a more inclusive list of species that exhibit this unique proteolysis, we analyzed the apparent molecular weight of the subunit α in other cell lines and subspecies (Figure 1B). Whole cell lysates were generated from *T. brucei* Lister 427 PF, *T. congolense* IL3000 PF, *T. b. gambiense* LiTat 1.3 BF, *T. brucei* EATRO 164 BF, *T. b. evansi* AnTat 3/3 BF, *L. major* Friedlin promastigotes (PM) and *L. mexicana* MNYC/BZ/62/M379 PM. The western blots from these samples were probed with a polyclonal antibody generated against the mature C-terminal α subunit of *T. brucei* (see methods and materials). A specific band was detected above the 40 kDa marker for each cell line, indicating conserved proteolysis of subunit α.

If the *T. brucei* subunit α did not undergo proteolytic processing, the UCSF ChimeraX visualization program predicts that the octapeptide would correspond to a loop that extends out from the yeast subunit α (Figure 1C) [44]. Therefore, we tested if this easily accessible loop from *S. cerevisiae* subunit α is also cleaved when expressed in *T. brucei.* A C-terminally V5-tagged yeast subunit α (Sc sub α FL WT V5) was expressed in *T. brucei* PF cells for 48 h, after which cytosolic and organellar fractions were analyzed by SDS-PAGE and western blotting. Blots probed with anti-V5 and compartment-specific markers (APRT for cytosol; mtHSP70 for mitochondrion) (Figure 1D) showed that Sc sub α FL WT V5 was efficiently targeted to mitochondria, where a predominant band of the expected full-length protein (58 kDa) is observed. These results suggest that euglenozoan subunit α proteolysis depends on specific sequence recognition sites.

The dual proteolytic cleavages that remove the subunit α octapeptide and generate separate N- and C-terminal polypeptides have so far been precisely defined only in *T. brucei*. However, the corresponding sequence (LFTKAAPLLDD) is also absent in the cryo-EM F_1_F_o_-ATP synthase dimer structure of the euglenozoan *Euglena gracilis*, a non-parasitic alga [19]. This indicates that subunit α octapeptide processing is conserved across Euglenozoa. Peptidase selectivity depends on the recognition of the specific chemistry or topography of the three to four amino acids before or after the cleavage site [45,46]. To identify a possible recognition sequence for the protease(s) involved in subunit α processing, we performed a Clustal Omega alignment of the subunit α genes from representative species within Euglenozoa (Figure 1E). The alignment revealed significant sequence conservation between most of the species in the region prior to the octapeptide. While the EV β strand identified in the *T. brucei* subunit α structure upstream of the octapeptide is not observed in the yeast or bovine structures [47,48], the position of the valine residue is consistently identified throughout eukaryotes as a hydrophobic aliphatic amino acid (i.e. valine, leucine or isoleucine) [43]. In contrast, the frequently conserved euglenozoan amino acids PVGL, located immediately N-terminal of the octapeptide, have strikingly different chemistries compared to the corresponding DGKG sequence often found throughout the other major eukaryotic groups [43]. Strikingly, there is very little conserved identity among the residues immediately downstream of the octapeptide. To better illustrate the chemistry of the amino acids conserved within and immediately surrounding the expelled octapeptide in euglenozoans, we generated a weblogo (Figure 1F) [49] from the alignment in Figure 1E, but without the yeast sequence. In general, the PVGL residues N-terminal of the first octapeptide cleavage site are predominantly hydrophobic, with the exception of the polar glycine. Within the octapeptide, the first amino acid C-terminal of the first cleavage site is usually the hydrophobic phenylalanine, leucine or isoleucine. This is followed by the conserved TRSR amino acids that contain alternating polar and basic side chains. Finally, the last two residues of the octapeptide are hydrophobic methionine or leucine. These bioinformatic analyses highlight potentially important residues that might be recognized by the proteases responsible for *T. brucei* subunit α proteolysis.

### C-terminal tagging of subunit α destabilizes the F_1_F_o_-ATP synthase complex

To determine which of these amino acids are required for subunit α proteolysis in vivo, we initially implemented a molecular tool to track ectopically expressed subunit α by tagging a wildtype subunit α with a C-terminal V5 epitope (Supplementary Figure S1A). If this proved successful, we could then express variants of subunit α that contained mutations around the proteolytic cleavage sites. However, tetracycline induction of the V5 tagged subunit α led to a moderate growth phenotype of the parasites (subunit α FL^WT^ V5) grown in glucose-rich SDM-79 medium (Supplementary Figure S1B). Western blot analyses of whole-cell lysates from this cell line reveal that subunit α FL^WT^ V5 underwent the expected proteolytic cleavage, with the V5 antibody predominantly recognizing a ∼53 kDa product (Supplementary Figure S1C). Subcellular fractionation further indicated that the processed C-terminal fragment was correctly targeted to the mitochondrion, as it was almost exclusively detected in the organellar fraction (Supplementary Figure S1C). Blue-native (BN) western blot analysis of isolated mitochondria demonstrated that the V5-tagged C-terminal polypeptide was incorporated into assembled F_1_-ATPase complexes as well as F_1_F_o_-ATP synthase monomers and oligomers (Supplementary Figure S1D, left panel). However, we consistently observed reduced levels of assembled F_1_- and F_1_F_o_-ATP synthase complexes containing subunit β upon the tetracycline induction of subunit α FL^WT^ V5 (Supplementary Figure S1D, right panel). To verify equal loading, mitochondrial material was denatured and probed with anti-mtHSP70, which revealed comparable protein levels between induced and non-induced cells (Supplementary Figure S1E).

In summary, it appears that the incorporation of the C-terminally tagged subunit α into the F_1_F_O_-ATP synthase complexes interferes with the overall stability of these rotary machines. Since the subunit α C-terminus ends with an α helix that folds back into the rest of subunit α [27], it is possible that the V5 tag creates steric hindrance problems during the assembly of the F_1_-ATPase. Due to the observed F_1_-ATPase complex instability and the identified N-terminal mitochondrial targeting peptide of subunit α [27], there are significant limitations to tagging the protein at either the N- or C-terminus. Thus, we opted for internal tagging using a FLAG epitope.

### Replacement of the octapeptide with a FLAG epitope generates an uncleaved subunit α that is incorporated into the F_1_F_O_ complex

We generated a cell line with a tetracycline-inducible ectopic copy of full-length subunit α in which the octapeptide has been replaced with the eight amino acid FLAG epitope (sub α FL^FLAG^, Figure 2A). Not only will this allow us to identify the ectopic mutant subunit α, but it can also be used to discern if the octapeptide contains amino acids essential for protease recognition. Importantly, while this epitope shares no similarity with the euglenozoan octapeptide sequence, the AI system AlphaFold predicts the FLAG epitope to form a loop between the mature N- and C-terminal subunit α polypeptides (Figure 2B) [50,51], similar to what is seen in the *S. cerevisiae* structure (compare Figure 1C) [48]. The induction of this subunit α FL^FLAG^ in the background of endogenous subunit α did not produce any growth defects even in low-glucose SDM-80 medium, where sufficient ATP production depends on oxidative phosphorylation (Figure 2C). Western blot analyses performed with a FLAG primary antibody revealed a predominate band that corresponded in size (∼61 kDa) to the uncleaved *T. brucei* subunit α (Figure 2D). This suggests that the proteolytic recognition sequence includes – or perhaps is even largely constrained to - the octapeptide. Furthermore, we only detected this subunit α FL^FLAG^ in the organellar subcellular fraction (Figure 2D), indicating that it was efficiently targeted to the mitochondrion.

**Figure 2.**
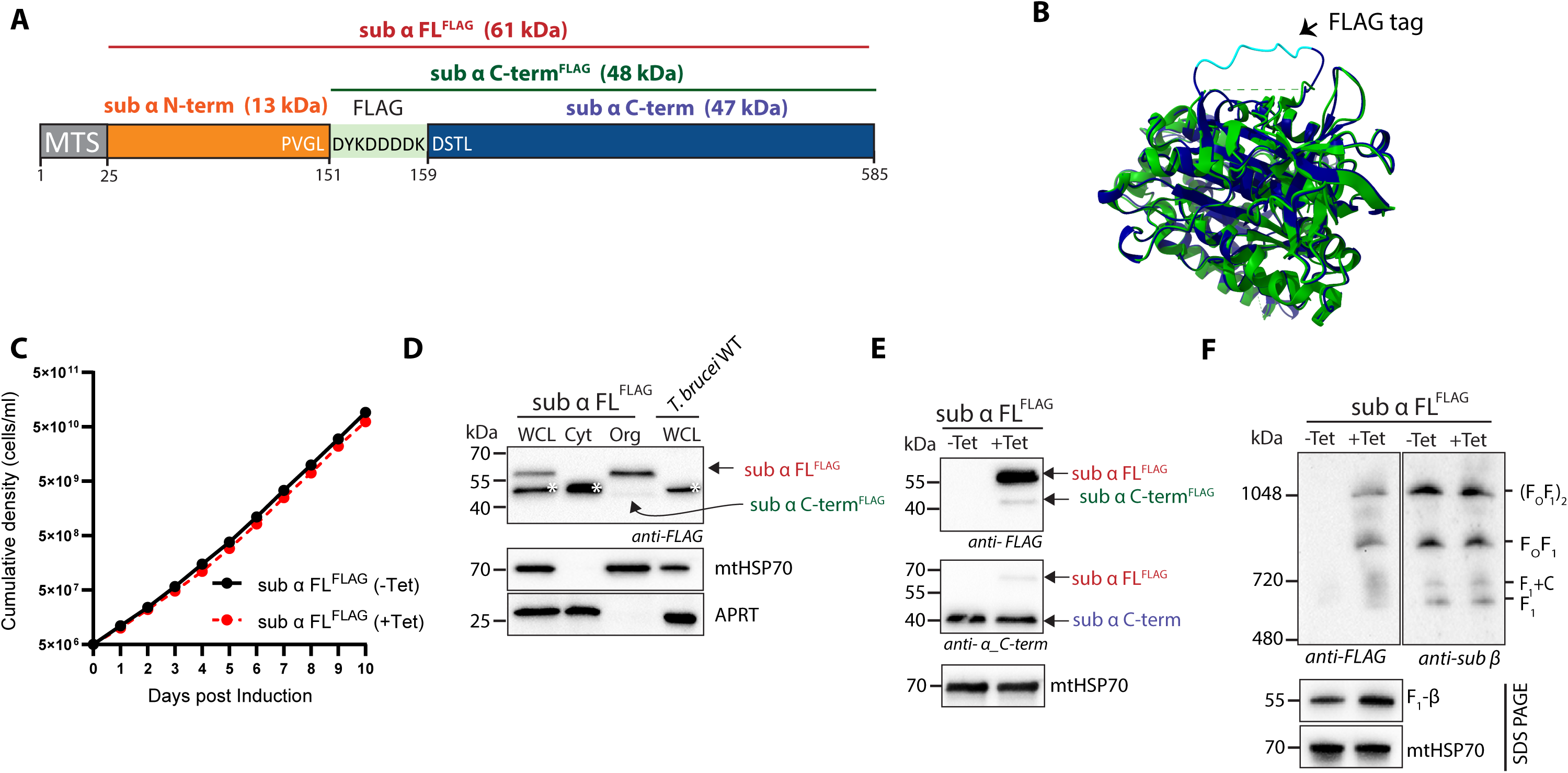
Octapeptide replacements for FLAG tag yields uncleaved subunit α, which incorporates into the F_1_F_o_-ATP synthase. **A.** Schematic of *T. brucei* subunit α with a FLAG tag replacing the octapeptide (sub α FL^FLAG^). **B.** Superimposition of the *T. brucei* subunit α FL^WT^ (6F5D, green) with the Alphafold predicted structure of subunit α FL^FLAG^ (blue). The RMSD between 517 pruned atom pairs is 0.424 Å. The FLAG tag (cyan) is predicted to form a flexible loop. **C.** Growth curve analysis of *T. brucei* cells induced to express sub α FL^FLAG^. Noninduced (-Tet) cells, black line; induced (+Tet) cells, red line. **D.** Immunoblots of whole cell lysates and subcellular fractions from subunit α FL^FLAG^ cells induced with tetracycline for 48 hours. Samples were probed with anti-FLAG, anti-mtHSP70 (mitochondrial marker) and anti-APRT (cytosolic marker). The relevant sizes of the protein marker are indicated on the left. A white asterisk indicates an unspecific band visualized in the cytosolic fraction with the anti-FLAG antibody. **E.** Western blot analysis of mitochondrial lysates from subunit α FL^FLAG^ noninduced (-Tet) and induced (+Tet) cells using anti-FLAG, anti-subunit α C-term and anti-mtHSP70 antibodies. The relevant sizes of the protein marker are indicated on the left. **F.** Representative immunoblots of mitochondrial lysates from subunit α FL^FLAG^ noninduced (-Tet) and induced (+Tet) cell lines resolved by BN-PAGE probed with antibodies against FLAG and subunit β. Positions of high molecular weight markers are shown. SDS PAGE analysis followed by western blot of the same samples probed with anti-HSP70 antibody served as a loading control.

A faint band around 48 kDa in the mitochondrial fraction, clearly distinct from the cross-reacting cytosolic protein (labeled with *), suggested the possibility of very minor processing of subunit α FL^FLAG^ resulting in a C-terminal fragment with the FLAG epitope at its N-terminal end (indicated as sub α C-term^FLAG^ in Figure 2D). To investigate this possibility further, we examined western blots containing proteins from enriched mitochondria of cells that were either induced or non-induced for subunit α FL^FLAG^ expression. We detected a very faint FLAG signal (∼48 kDa) only in fractions from induced cells, again suggesting a partial proteolysis that resulted in a C-terminal subunit α that has retained the FLAG epitope (Figure 2E). This implies that sequence features located immediately upstream of the first octapeptide cleavage site might be able to recruit the proteolytic activity, albeit inefficiently. When these same western blots were probed with the subunit α C-terminal polyclonal antibody, it became apparent that the ectopic subunit α FL^FLAG^ expression was consistently low compared to the endogenous subunit α. Importantly, at least some of the uncleaved subunit α FL^FLAG^ is incorporated into F_1_- and F_1_F_O_-ATP synthase complexes, as observed in western blots of BN gels probed with the FLAG antibody (Figure 2F). Probing of the same blots with the subunit β antibody suggested that the total amount of ATP synthase complexes was not diminished by subunit α FL^FLAG^ expression. Thus, replacing the octapeptide sequence with an unrelated sequence creates a subunit α mutant that is largely resistant to proteolysis, albeit with reduced levels of expression. The uncleaved subunit α FL^FLAG^ is incorporated into apparently complete F_1_F_O_-ATP synthase complexes, suggesting that proteolytic processing of this subunit is not a pre-requisite for ATP synthase assembly.

### Subunit α proteolysis is not essential for PF *T. brucei* grown in culture

To assess if F_1_F_o_-ATP synthase with an uncleaved α subunit is fully functional, we asked whether expression of either a recoded ectopic wildtype α (sub α RNAi rec FL^WT^) or a recoded subunit α where the octapeptide was replaced with an uncleavable linker sequence (sub α RNAi rec FL^PAS^) would rescue the growth defect of a subunit α RNAi cell line (sub α RNAi) (Figure 3A). For these experiments, instead of the FLAG epitope, we chose the semiflexible PAS-motif linker, which consists of randomly composed proline, alanine and serine residues. This linker is capable of expansion, while also being resistant to aggregation and proteolysis [52,53]. When the subunit α FL^PAS^ structure predicted by Alphafold is superimposed on the subunit α FL^WT^ crystal structure, the PAS linker is depicted as an exposed loop (Figure 3B). Exchanging the FLAG epitope with a flexible PAS linker to bridge the N- and C-terminal subunit α peptides was also expected to increase ectopic expression levels by adding some stability to the mutant subunit α.

**Figure 3.**
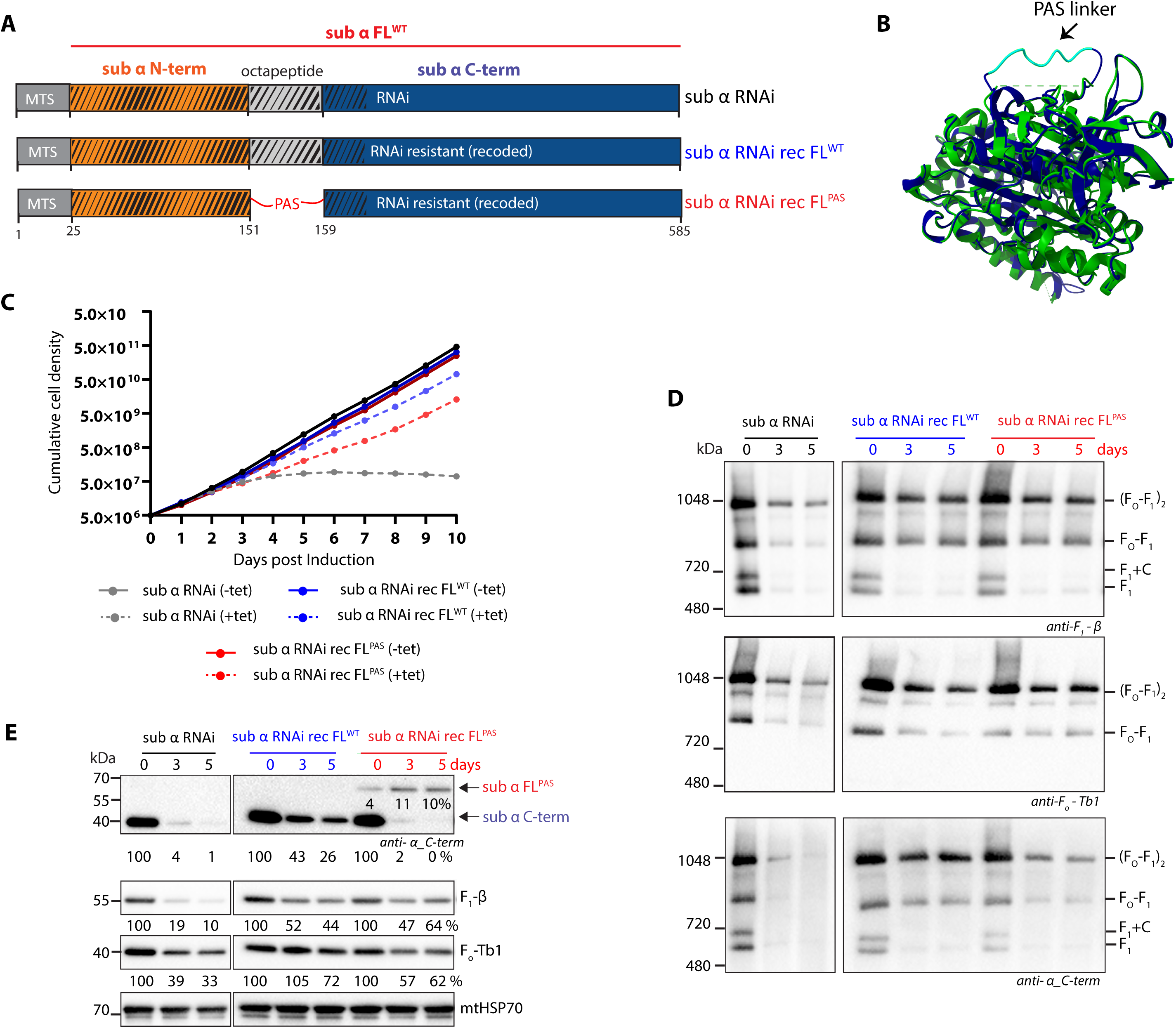
Replacing the octapeptide with a flexible PAS linker results in an uncleaved subunit α that rescues the growth phenotype observed upon subunit α RNAi induction. **A.** Schematic of *T. brucei* subunit α FL^WT^ with the region targeted by RNAi (hatched, sub α RNAi), its RNAi-resistant version containing a recoded (rec) subunit α gene (sub α RNAi rec FL^WT^) and recoded subunit α containing a PAS linker instead of the octapeptide (sub α RNAi rec FL^PAS^). **B.** Superimposition of *T. brucei* subunit α FL^WT^ (6F5D, green) with the Alphafold predicted structure for subunit α FL^PAS^. The RMSD between 518 pruned atom pairs is 0.417 Å. The PAS linker as a flexible loop is shown in cyan. **C**. Growth curves of subunit α RNAi, sub α RNAi rec FL^WT^ sub α RNAi rec FL^PAS^ noninduced (-Tet) and RNAi induced (+Tet). **D.** Representative immunoblots of mitochondrial lysates from the various subunit α RNAi cell lines that were either noninduced (-Tet) or induced for 3 and 5 days (+Tet) resolved by BN-PAGE probed with antibodies against F_1_-ATPase subunits α C-term and subunit β, and the F_o_-moiety subunit ATPTb1. Positions of high molecular weight marker are shown. The numbers beneath the blot represent the abundance of immunodetected proteins expressed as a percentage of the noninduced sample after normalizing to the signal intensity of the mtHsp70 probing (loading control). **E.** Mitochondrial lysates from **D** were separated on SDS PAGE and probed with antibodies against subunit α C-term, subunit β and ATPTb1. Immunoblot with anti-mtHSP70 served as a loading control.

We first measured the growth rate of the three cell lines in the presence and absence of the RNAi inducer tetracycline (Figure 3C). Compared to the growth defect produced when endogenous subunit α is depleted by RNAi (sub α RNAi +Tet), the constitutive expression of subunit α FL^WT^ almost completely rescued the RNAi growth defect. Constitutive expression of subunit α FL^PAS^ resulted in a clear, although partial, rescue of the growth defect. Next, we fractionated enriched mitochondrial fractions by BN gel electrophoresis and probed western blots of these gels with specific antibodies to determine the amount of F_1_F_O_-ATP synthase complexes that were present in each cell line (Figure 3D). We used antibodies recognizing either the F_1_ subunit α C-terminal peptide, F_1_ subunit β, or the F_o_ subunit Tb1. Induction of subunit α RNAi in the parental cell line for 3 and 5 days resulted in a progressive loss of monomeric and dimeric F_1_F_o_-ATP synthase complexes, as well as of the partial F_1_ and ‘F_1_+C’ complexes (Figure 3D, left panel; ‘F_1_+C’ refers to a partial complex composed of F_1_ and the subunit *c* ring) [54], consistent with what has been observed in earlier studies [20,55]. The amount of F_1_F_o_-ATP synthase complexes detected after RNAi induction in either the sub α RNAi rec FL^WT^ or sub α RNAi rec FL^PAS^ samples was relatively equivalent when immunolabelled with the subunit β or Tb1 antibodies. While these levels of the enzyme were less than what was observed in the noninduced day 0 samples, they were more abundant than in the parental sub α RNAi induced samples, indicating assembly of the ectopically expressed subunit α into partial and complete F_1_F_o_ complexes. However, probing the blots with the subunit α C-terminal antibody indicated that there might be less enzyme in the sub α RNAi rec FL^PAS^ than the sub α RNAi rec FL^WT^ samples upon RNAi induction. This effect may reflect a reduced antibody affinity for the C-terminal region of the unprocessed subunit α FL^PAS^. We also analyzed these mitochondrially enriched samples by resolving them on denaturing SDS-PAGE and probing blots with the same antibodies (Figure 3E). When probed with the subunit α C-terminal antibody, we observed a band of ∼60 kDa only in the sub α FL^PAS^ samples, which increased in intensity over the course of the RNAi induction. This observation is consistent with production of a proteolysis-resistant subunit α that, in the absence of the endogenous subunit, becomes stabilized by incorporation into the F_1_F_O_ complex. Amounts of the mature subunit α C-terminal polypeptide were almost undetectable in these samples. While the levels of subunit α FL^PAS^ were reduced compared to (processed) subunit α FL^WT^, they were clearly higher than in the parental sub α RNAi induced samples. Finally, while the levels of F_1_-ATPase subunit β are again similar between the two rescue cell lines, the amount of Tb1 seems to be slightly lower in the sub α RNAi rec FL^PAS^. In summary, constitutive expression of an RNAi-resistant subunit α where the octapeptide has been replaced with a PAS linker results in production of proteolytically resistant protein that, in the absence of the endogenous protein, gets incorporated into fully assemble F_1_F_o_-ATP synthase complexes. The growth defect of subunit α RNAi is rescued in this cell line, but only partially, which could be due to suboptimal expression levels and/or impaired activity of an ATP synthase complex with an unprocessed α subunit.

### Incorporation of proteolytically resistant subunit α FL^PAS^ into the F_1_F_O_ complex does not affect ATP synthase activities

To determine if F_1_F_o_-ATP synthase complexes containing uncleaved subunit α FL^PAS^ have altered rates of catalytic activity, we analyzed the ATP production [56] capabilities of digitonin permeabilized sub α RNAi rec FL^WT^ and sub α RNAi rec FL^PAS^ cells after 5 days of RNAi induction (Figure 4A). The amount of ATP produced ‘in-organello’ via oxidative phosphorylation can be determined by incubation with ADP and glycerol-3-phosphate (Gly-3-P) as substrates. The Gly-3-P is oxidized by the mitochondrial Gly-3-P dehydrogenase, which then passes the newly obtained electrons through respiratory complexes III and IV via ubiquinol and cytochrome c, respectively. The F_1_F_o_-ATP synthase then converts the supplied ADP into ATP using the membrane potential generated by the proton-pumping of respiratory complexes III and IV. Both the sub α RNAi rec FL^WT^ and sub α RNAi rec FL^PAS^ produced similar amounts of ATP. The decreased ATP produced when the respiratory complex III inhibitor antimycin A (ANT) or respiratory complex IV inhibitor potassium cyanide (KCN) are added indicates that the observed ATP production is primarily due to oxidative phosphorylation.

**Figure 4.**
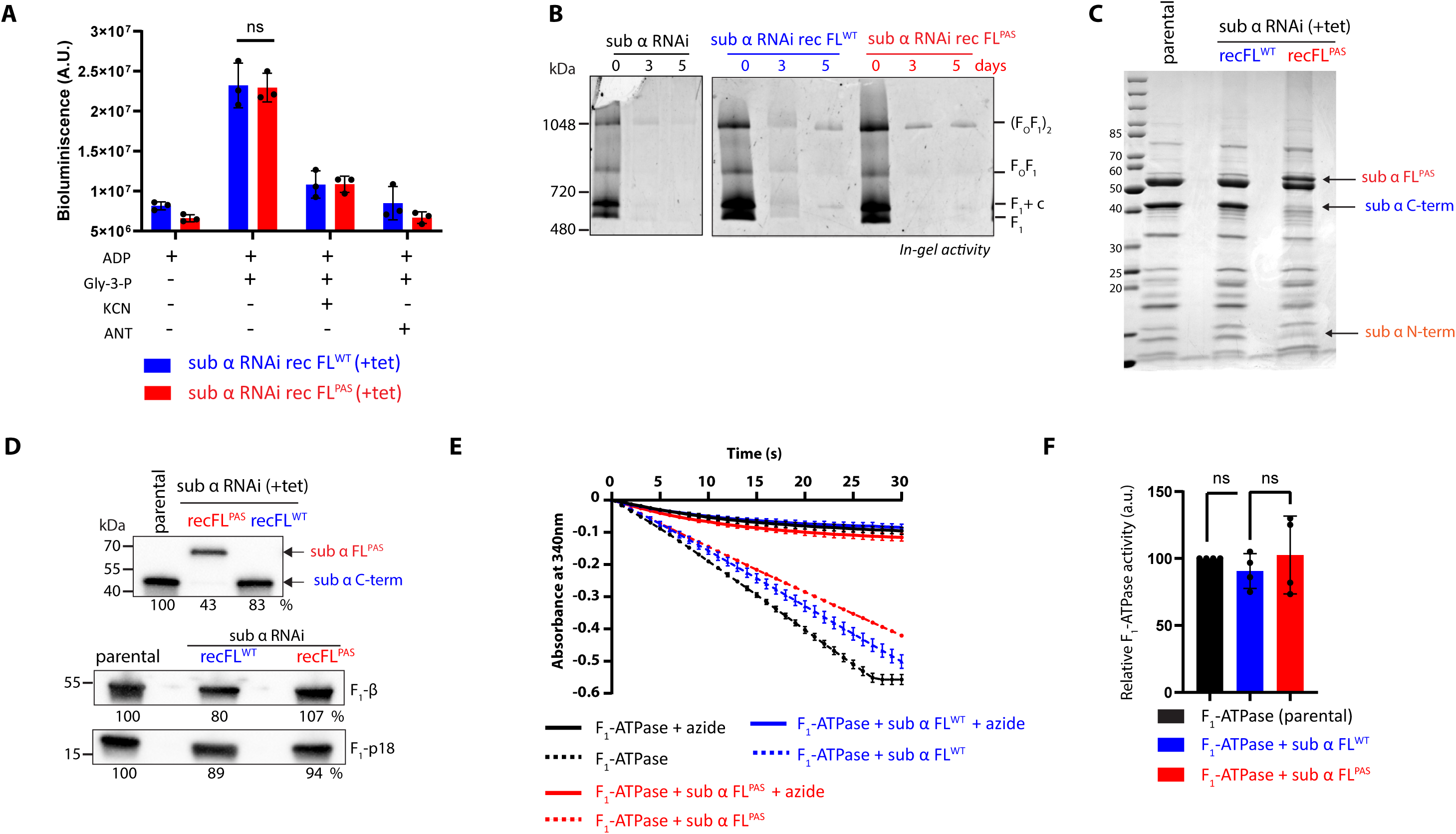
Uncleaved version of subunit α containing flexible PAS linker rescues RNAi depleted F_1_F_O_-ATP synthase activities. **A.** ATP production in digitonin-permeabilized subunit α RNAi rec FL^WT^ and subunit α RNAi rec FL^PAS^ induce for 5 days. The graphs show individual values of three biological replicates, their means and standard deviations. The substrate (Gly-3-P, D,L-glycerol-3-phosphate), ADP and inhibitors (KCN, potassium cyanide, ANT, antimycin) were added as indicated. **B.** Blue native electrophoresis of dodecyl maltoside-lysed mitochondrial lysates (40 µg) from the RNAi non-induced and the 3 and 5 day RNAi-induction of the indicated cell lines werefollowed by in-gel staining of ATPase activity. **C.** Purified F_1_-ATPase (10 µg) from the parental 29-13 cell line and subunit α RNAi rec FL^WT^ and subunit α RNAi rec FL^PAS^ was fractionated on SDS PAGE gel followed by Coomassie staining. Positions of full-length (FL) α and its C- and N-terminal polypeptides are labeled by arrows. **D.** Immunoblot of F_1_-ATPase samples (3 µg) using antibody against subunit α C-term, subunit β and p18. The numbers beneath the blot represent the abundance of immunodetected proteins expressed as a percentage of the parental sample. **E**. ATPase activity assay using chloroform-extracted F_1_-ATPase particles from *T. brucei* parental 29-13, subunit α RNAi rec FL^WT^ and subunit α RNAi rec FL^PAS^ induce for 5 days. As a control for specificity, the samples were also treated with 2 mM azide. The graph is a representative experiment showing individual values of three technical replicates for each of the cell lines (n=3, ± s.d.) **F.** ATPase specific activity was quantified using the Beer-Lambert law and then normalized based on protein levels determined by western blot analysis (**D**) and Coomassie-stained gel (**C**). The activity of the F_1_-ATPase from the parental 29-13 strain was set to 100%.

Next, we compared F_1_-mediated ATP hydrolysis activities between RNAi-induced parental, sub α RNAi rec FL^PAS^ and sub α RNAi FL^WT^ cell lines. First, we used an in-gel activity assay (Figure 4B). After 5 days of subunit α RNAi induction, we observed similar amounts of lead phosphate precipitation in both sub α RNAi rec FL^WT^ and sub α RNAi rec FL^PAS^, indicating comparable activity of wild-type complexes and complexes with uncleaved α subunit. In both cases, ATP hydrolysis staining was more intense than the parental sub α RNAi induced samples, consistent with partial rescue of the enzyme activity.

Both the in-organello ATP production assay and the in-gel ATP hydrolysis assay are endpoint assays that might obscure potential differences in the rates of activity. Therefore, we also isolated F_1_-ATPase complexes by using chloroform extraction to release the enzymatic domain from the mitochondrial inner membrane [43,57,58]. With only a few impurities remaining from this fast purification of the enzyme, each of the F_1_-ATPase subunits can be identified when resolved by SDS-PAGE and then stained with coomassie (Figure 4C). The two most prominent bands observed for the parental and the sub α rec FL^WT^ cell line represent the F_1_-ATP synthase subunit β (∼54 kDa) and the processed subunit α C-terminal polypeptide (∼47 kDa). For the sub α RNAi rec FL^PAS^ cell line, the expected shift for uncleaved subunit α to ∼61 kDa was observed. This was further corroborated by western blot analyses of these samples (Figure 4D). The top blot probed with the subunit α C-terminal antibody reiterates that a processed subunit α C-terminal peptide was almost undetectable in the sub α RNAi rec FL^PAS^ cells. These same F_1_-ATPase purifications were also probed with antibodies specific for the F_1_-ATPase subunits β and p18. These immunoblots were then analyzed by scanning densitometry to quantify the relative abundance of the purified F_1_-ATPase from each cell line. Again, we observed reduced levels of subunit α in sub α RNAi rec FL ^PAS^ samples, while the other two F_1_-ATPase subunits showed comparable abundance across the different cell lines.

We then performed a Pullman assay [59] to measure the kinetic rate of ATP hydrolysis for the isolated F_1_-ATPase complexes (Figure 4 E). The ATP hydrolysis rates were measured for equal amounts of F_1_-ATPase from various cell lines (Supplementary Table S4). These rates were then normalized based on the average protein levels determined either by western blot (Figure 4D) or coomassie stained gels (Figure 4C) and compared to the ATP hydrolysis rates determined for the wildtype enzyme (Figure 4F). This analysis found no significant differences between the ATP hydrolytic rates of F_1_-ATPase complexes containing either a proteolytically cleaved or uncleaved subunit α.

### FRET peptide assay indicates significant *T. brucei* subunit α proteolysis in the cytosol

The proteolytic cleavage events could conceivably take place in the cytosol, before mitochondrial import (with separate import of the two cleavage products), or after mitochondrial import, before or after incorporation into the ATP synthase complex. To investigate these possibilities, we devised a Förster resonance energy transfer (FRET) reporter system that allowed us to identify when a peptide corresponding to either the upstream (PVGL|VTRS) or downstream (RRLL|DSTL) cleavage sites of the octapeptide underwent proteolysis. The eight amino acid FRET peptides were designed with an N-terminal quencher (DABCYL) and a C-terminal fluorescence donor (EDANS), which results in a quenched fluorescence until the donor is released from the quencher by proteolysis. *T. brucei* cells were fractionated into subcellular material by digitonin latency [60,61]. Digitonin preferentially interacts with the cholesterol in membranes, which leads to the formation of pores. The amount of cholesterol in various organellar membranes varies, so we can use this property to enrich specific organelles [62–64]. A pilot digitonin analysis revealed that the cytosolic protein APRT was released from the whole cell from as little as 0.02 mg digitonin/mg total cellular protein (Supplementary Figure S2). Whereas the glycosomal markers hypoxanthine-guanine-xanthine phosphoribosyltransferase (XPRT) and hexokinase were released from 0.06 mg digitonin/mg of total cellular protein. Mitochondrial proteins mtHSP70 and succinyl-CoA ligase subunit β (SCS) were released at 0.16 mg digitonin/mg of total cellular protein. With this information, we devised the digitonin latency scheme presented in Figure 5A to separate the cytosolic and mitochondrial fractions from the same starting pool of cells. Western blot analyses of each generated subcellular fraction were performed with markers for the cytosol, glycosomes and mitochondria (Figure 5B). These FRET peptides were then incubated with equal amounts of protein from the cytosolic or mitochondrial fractions (Figure 5C, D). The control experiments, which included only the FRET peptide or just the *T. brucei* subcellular fractions, resulted in minor amounts of background fluorescence. While both FRET peptides incubated with the mitochondrial fraction gave some steady state level of fluorescence, this rate was much less than what we observed when the FRET peptides were incubated with the cytosolic fraction. This suggests that the protease(s) responsible for *T. brucei* subunit α proteolysis may reside in the cytosol.

**Figure 5.**
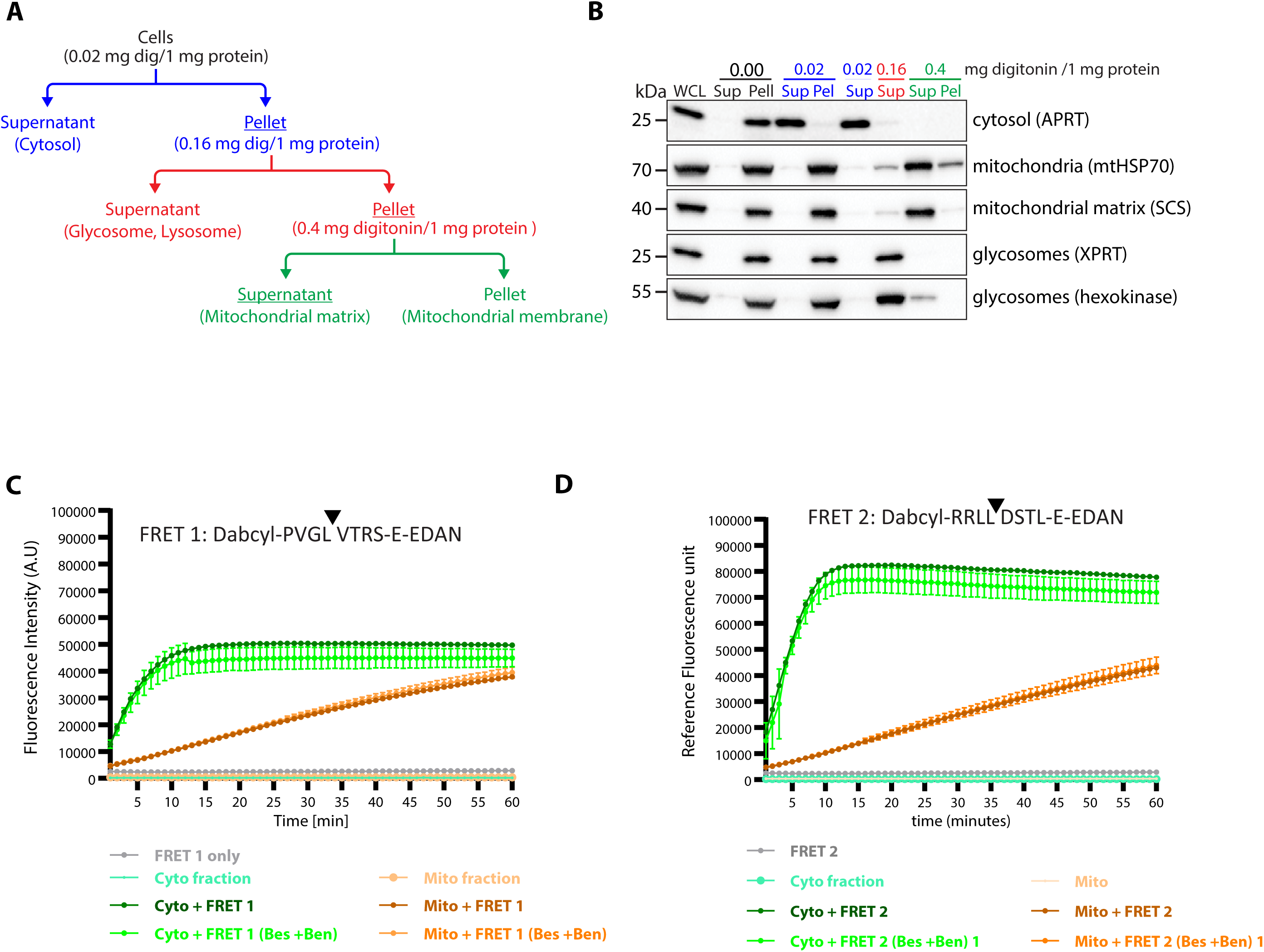
Cytosolic fraction produces significant FRET activity for the two peptides mimicking subunit α octapeptide cleavage sites. **A.** Schematic for digitonin latency protocol to obtain highly enriched cytosolic and mitochondrial fractions from a single population of cells. **B.** Immunoblot analysis of digitonin-obtained fractions using specific antibodies marking different cellular compartments. **C.** Fluorescence activity of FRET peptide 1 and 2 upon addition of cytosol and mitochondrial matrix fractions. Protease inhibitors bestatin (bes) and benzylsuccinic (bez) acid were used to inhibit non-specific protease activity.

### Individual ectopic expression of either the *T. brucei* subunit α N-terminal or C-terminal peptides results in mitochondrial localization

Localisation of the proteolytic cleavage activities in the cytosol, prior to mitochondrial import, would imply that the cleavage products can be imported individually, and that the C-terminal polypeptide must therefore contain a separate import signal. In order to investigate this hypothesis, we expressed in PF *T. brucei* either the full-length subunit α coding sequence, the N-terminal peptide with the defined mitochondrial targeting sequence or the mature C-terminal peptide. A C-terminal V5 tag was added to distinguish the ectopic gene products from endogenous subunit α polypeptides. This resulted in the following cell lines: sub α FL^WT^ V5 (*T. brucei* subunit α amino acids 1-584), sub α N-term V5 (amino acids 1-151) and sub α C-term V5 (amino acids 160-584). The subcellular targeting of these subunit α peptides was first analyzed by immunofluorescence microscopy using a primary antibody recognizing the V5 epitope (Figure 6A). As expected, sub α N-term V5, like the control sub α FL^WT^ V5, was efficiently targeted to the highly branched mitochondrion. Interestingly, the subunit α V5 tagged C-terminal peptide was also targeted to the mitochondrion, suggesting the presence of an internal mitochondrial targeting signal. This immunofluorescence data is further corroborated by the western blot analyses of the cytosolic and organellar subcellular fractions from each of these cell lines (Figure 6B). While there is a very minor proportion of subunit α V5 tagged C-terminal peptide that is retained in the cytosolic fraction, this is also detected in the control sub α FL^WT^ V5. Most likely this is the result of high levels of ectopic protein expression induced with 1 ug tetracycline overwhelming the mitochondrial import machinery. Together, these results demonstrate that if subunit α proteolysis occurs in the cytosol, the C-terminal polypeptide likely possesses an internal mitochondrial targeting signal that enables its independent mitochondrial import.

**Figure 6.**
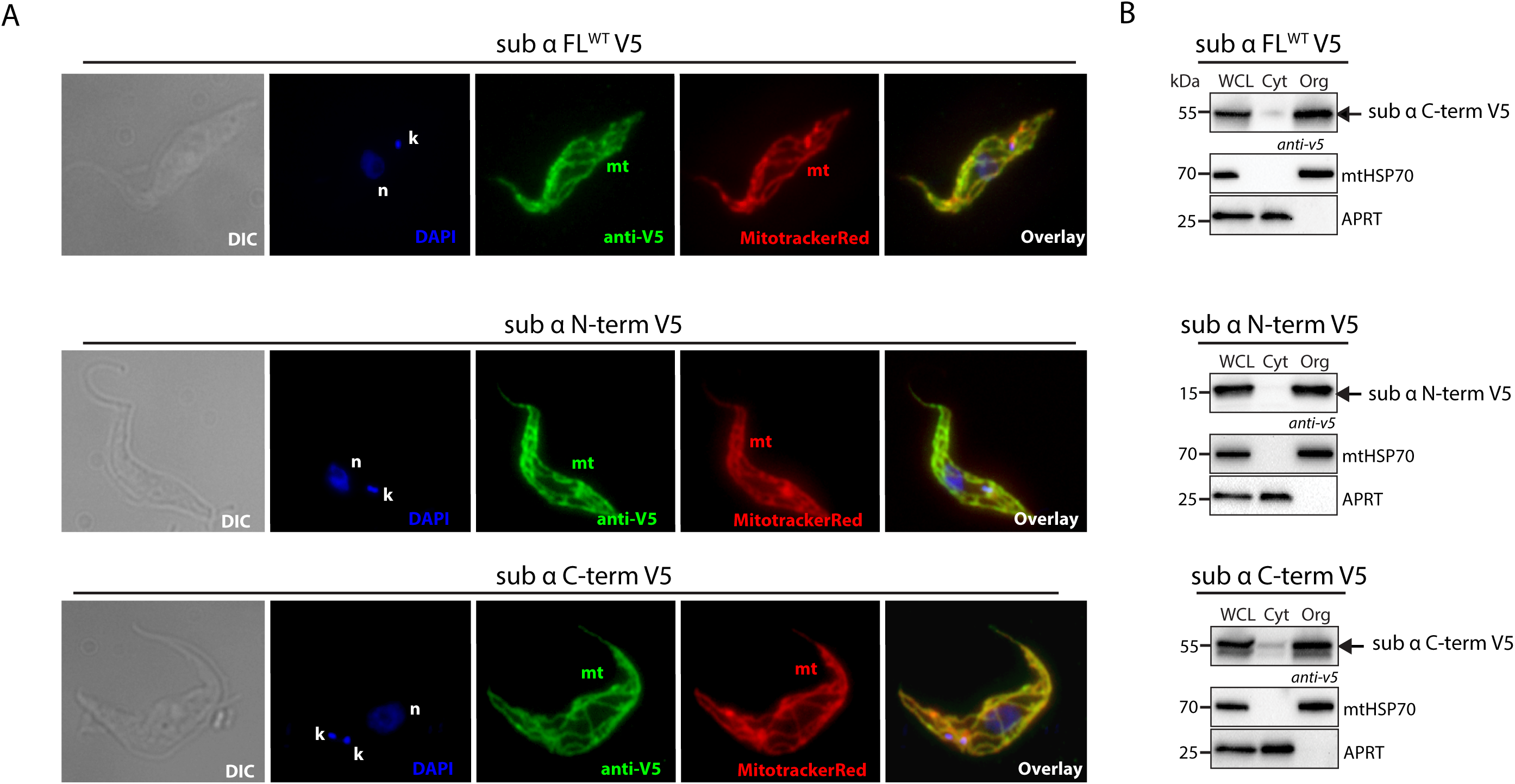
Subunit α N-terminal and C-terminal polypeptides are imported into the *T. brucei* mitochondrion. **A.** Immunofluorescence assay of tetracycline induced *T. brucei* cell lines expressing either subunit α FL^WT^, subunit α N-terminus or subunit α C-terminus tagged with the V5 epitope. Fluorescence was observed from a Alexa Fluor 488-conjugated secondary antibody recognizing the primary anti-V5 antibody. The DNA content and the single reticulated mitochondrion were visualized using DAPI (4,6-diamidino-2-phenylindole) and Mitotracker Red CMXRos staining, respectively. The overall cell morphology is depicted in the differential interference contrast (DIC) microscopy images. **B.** Immunoblots of whole cell lysates (WCL) and subcellular fractions comprised of cytosolic (Cyt) or organellar (Org) material from subunit α FL^WT^ V5, subunit α C-term V5 and subunit α N-term V5 cell lines induced with tetracycline for 48 hours. Samples were probed with anti-V5, anti-mtHSP70 (mitochondrial marker) and anti-APRT (cytosolic marker). The relevant sizes of the protein marker are indicated on the left.

## CONCLUSIONS

Since the mitochondrial F_1_-ATPase in eukaryotes was long believed to be homogeneous in structure and composition our work set out to determine whether *T. brucei* subunit α proteolysis is biologically significant. Sequence alignments of subunit α from 14 euglenozoan species revealed conserved chemical features within and immediately upstream of the octapeptide cleavage site, which differ markedly from the corresponding region in *S. cerevisiae* and other eukaryotes. Despite overall structural similarity between the *T. brucei* and yeast proteins, the parasite was unable to cleave the heterologous yeast subunit α, underscoring the importance of these lineage-specific residues for proteolysis. By replacing the octapeptide with either a FLAG epitope or a PAS linker, we generated proteolysis-resistant subunit α mutants, demonstrating for the first time that some of the key recognition elements are contained within the octapeptide itself. These cleavage-resistant variants were incorporated into functional F_1_F_o_-ATP synthase complexes. In vitro biochemical assays measuring ATP production and ATP hydrolysis from complexes either resolved on native gels or biochemically purified, suggest that *T. brucei* subunit α proteolysis is not essential for either the enzymatic activity of the rotary machine or for the growth of the cultured procyclic parasites.

Nevertheless, important questions remain regarding the physiological role of this unusual processing event during the parasite’s complex life cycle. Future studies should extend these analyses to the bloodstream form of *T. brucei*, which relies exclusively on the hydrolytic activity of the complex. The physiological relevance of the cleavage could also be examined during the in vitro differentiation of bloodstream forms into procyclic forms. This system would allow testing whether subunit α proteolysis contributes to the activity switch from ATP hydrolysis to ATP synthesis that underlies parasite development and adaptation to new environments. In addition, the identity of the responsible protease(s) remains unknown. Determining whether the activity arises from a conserved eukaryotic protease that recognizes the variant euglenozoan octapeptide sequence, or from a unique trypanosomatid enzyme, will be critical for understanding the evolutionary significance of this process. The FRET-based assays developed here provide a valuable tool to address these questions.

Taken together, our findings reveal that subunit α proteolysis is a conserved, sequence-specific event in euglenozoans and it highlights an alternative pathway of ATP synthase biogenesis. By separating a single subunit into independently imported N- and C-terminal fragments, this mechanism expands the repertoire of strategies by which eukaryotes assemble their molecular machines. This distinctive feature may contribute to the remarkable plasticity of mitochondrial structure and function in trypanosomes, organisms that must rewire their energy metabolism during transitions between host environments. More broadly, the work illustrates how lineage-specific protein processing can diversify the architecture of a universally conserved enzyme, raising the possibility that similar hidden variations remain to be discovered in other protist lineages.

## METHODOLOGY

### Trypanosomatid parasites cell cultures

The procyclic form (PF) *T. brucei* Lister 427 strain was grown at 27°C in high-glucose SDM-79 or low-glucose SDM-80 medium with 10% fetal bovine serum (FBS) and 7.5 mg/ml hemin. The first medium supplies nutrients for glycolysis to support substrate phosphorylation which the latter one supplies amino acids, as an energy source produced by oxidative phosphorylation. PF 29-13 Lister 427 and the *T. brucei* SMOX Lister 427 strains expressing T7 RNA polymerase and tetracycline repressor were grown with 15μg/ml G418 and 25 μg/ml hygromycin B Gold or with 0.1 μg/ml puromycin, respectively. PF *T. congolense* IL3000 was maintained in TcPCF-3 media [65] with 20% FBS at 27°C.

The bloodstream form (BF) *T. brucei* Lister 427, acriflavine-induced dyskinetoplastic *T. brucei* EATRO 164 [66] and dyskinetoplastic *T. b. evansi* AnTat 3/3 [36] were grown at 37°C and 5% CO_2_ in HMI-11 medium with 10% FBS. *T. b. gambiense* BF strain was maintained in HMI-11 medium with 20% FBS. Promastigote *L. major* Friedlin and *L. mexicana* MNYC/BZ/62/M379 were grown at 25°C in M199 medium with 10% FBS.

### Plasmid construction and generation of transgenic *T. brucei* cell lines

To generate overexpression transgenic cell lines subunit α FL WT V5, subunit α N-term V5, subunit α C-term V5 and *S. cerevisiae* subunit α V5 (Table S1), coding sequences (cds) were amplified by PCR from PF *T. brucei* Lister 427 strain genomic DNA (gDNA) and *S. cerevisiae* BY4742 strain, respectively, using primers with restriction sites (Table S1 and S2). The PCR products were ligated to p2T7-3V5 plasmid [67]. The subunit α FL^FLAG^ mutant, subunit α RNAi rec FL^WT^ and subunit α RNAi rec FL^PAS^ were generated by a fusion PCR technique. Overlapping PCR fragments A and B were amplified using designated primers and cDNA sources as listed in Table S1. Next, fragment C was produced by combining A and B with specific primers including restriction sites (Table S2). The product for subunit α FL^FLAG^ was ligated to pLEW79 [68], while the two RNAi recoded fragments were cloned to pHD1344 [69]. All plasmids were verified by sequencing, linearized with NotI and transfected into different parental cell lines to integrate into specific genome locations (Table S1). Transgenic *T. brucei* cells were selected using vector-specific antibiotic resistance: puromycin (0.1 μg/ml) for pT7-3V5-PAC and pHD1344, and phleomycin (2.5 μg/ml) for pLEW79.

For growth curve analysis, *T. brucei* cell lines were induced with 1 μg/ml tetracycline for 7-10 days. Cell density was measured using a Beckman Z2 Cell Counter and maintained in their exponential mid-log growth phase at 5×10^6^ cells/ml. Exponential growth curves were prepared using GraphPad prism.

### Subunit α C- terminal peptide expression, purification and antibody preparation

The subunit α part of the gene corresponding to the C-term (without the octapeptide) was PCR-amplified from *T. brucei* Lister 427 gDNA and cloned into the pSKB3 plasmid for expression with an N-terminal 6xHis tag. The plasmid was transformed into *E. coli* Rosetta DE3 cells, which were grown in LB medium and induced with 1mM IPTG. After cell lysis with lysozyme (10mg/ml), the pellet was resuspended in the buffer containing 1.5% sarkozyl and the resulting supernatant was loaded on a HisTrap nickel affinity column using AKTA Prime chromatography system. The recombinant protein was eluted with imidazole and dialyzed. The final protein was concentrated, quantified, and submitted for antibody preparation at Davids Biotechnologies, Germany.

### SDS-PAGE

*T. brucei* cells were harvested and washed with PBS by centrifugation at 1300xg for 10 min at 4°C. The cell pellet was resuspended in PBS and 3xLaemmli buffer (150 mM Tris pH-6.8, 300 mM 1,4 dithiothreitol, 6%(w/v) SDS, 30% (w/v) glycerol, 0.02% (w/v) bromophenol blue) to obtain 30 µl of lysates representing material from 1×10^7^ parasites. Samples were denatured at 97°C for 10 min. For western blot analysis, 30 µl of sample was separated on pre-cast 4-20% gradient Tris-glycine polyacrylamide gels. Proteins were transferred onto a PVDF membrane, followed by incubation with an appropriate primary antibody. After washing, the membranes were probed with a secondary HRP-conjugated antibody (Table S3). The Clarity Western ECL substrate was then applied, and protein signals were visualized using a ChemiDoc MP Imaging System (Bio-Rad). The densitometric analysis of the bands was carried out using the ImageLab software by relating the signal intensity from the lanes corresponding to RNAi-induced cells to that of the lane pertinent to control cells. The percentage of downregulation relative to the control sample was then normalized to the corresponding signal intensity of the bands from the blots probed with mtHSP70 (loading control).

The dilutions of the primary antibodies, as well as the TriTrypDB GeneID of the recognized antigens, are listed here: mAb anti-v5 epitope tag (1:2000, Invitrogen), mAB anti-FLAG epitope (1:2000, Sigma), mAb anti-mitochondrial HSP70 (1:5000; TriTrypDB Gene ID: Tb927.6.3740, Tb927.6.3750 and Tb927.6.3800; 72 kDa), polyclonal antibodies (pAB) recognizing ATP synthase subunits β (1:2000; TriTrypDB Gene ID Tb927.3.1380; 54 kDa), N- and C-terminal subunit alpha (1:100 and 1:500, respectively; TriTrypDB Gene ID Tb927.7.7420/7430), p18 (1:1000; TriTrypDB Gene ID Tb927.5.1710; 18 kDa), Tb1 (1:1000; TriTrypDB Gene ID Tb927.10.520, 47 kDa), APRT (1:250, TriTrypDB Gene ID: Tb927.7.1780 and Tb927.7.1790, 28 kDa), succinyl-CoA ligase [GDP-forming] β-chain (SCS, 1:1000, TriTrypDB GeneID: Tb927.10.7410; 42 kDa), hypoxanthine-guanine-xanthine phosphoribosyltransferase (XPRT, 1:500; TriTrypDB Gene ID: Tb927.10.1390, 27 kDa) and hexokinase (1:500; TriTrypDB Gene ID: Tb927.10.2010, 57 kDa) (Table S3).

### F_1_-ATPase particle purification

Mitochondria from 2-3×10^10^ cells were isolated by hypotonic lysis and Percoll gradient centrifugation [70]. Mitochondria were the resuspended in buffer and adjusted to a protein concentration of 16 mg/mL. F_1_-ATPase was extracted as published earlier [57,71]. Briefly, samples were sonicated and subjected to overnight centrifugation to collect small membrane particles (SMPs). SMPs were then resuspended, and chloroform was added to release F_1_-ATPase. After centrifugation, the aqueous phase containing F_1_-ATPase was collected.

### BN-PAGE western blot and in-gel activity of F_1_F_o_-ATPase

2.5×10^8^ cell equivalents of hypotonically isolated mitochondrial vesicles were solubilized with 2% (w/v) DDM for 1 hour on ice. 6 µg of total protein was resolved on a native pre-cast PAGE 3– 12% Bis-Tris gel and transferred onto a PVDF membrane [72]. The membranes were probed with antibodies as previously outlined. For in-gel activity of F_1_F_o_-ATP synthase, 40 µg of mitochondrial protein lysate was resolved on the native gel. The F_1_ and F_1_F_o_-mediated ATP hydrolytic activities were visualized by incubating the gels in 40 ml of buffer consisting of 35 mM Tris-HCl pH-8.0, 19 mM MgSO4, 270 mM glycine, 0.3% (w/v) PbNO3 and 11 mM ATP overnight at room temperature [73]. As enzymes in the gel hydrolyze the added ATP into ADP and inorganic phosphate, the phosphate interacts with the lead and forms an opaque precipitate. The gel was fixed with 30% methanol.

### ATP hydrolysis assay

The ATP hydrolytic activity of purified F_1_-ATPase was measured spectrophotometrically using an ATP regenerating assay [59]. This assay is an enzyme coupled reaction that regenerates ATP as the substrates phosphoenolpyruvate (PEP) and ADP are converted first to pyruvate and then lactate. The latter step involves the oxidation of NADH to NAD+, which can be measured spectrophotometrically over time at 340 nm. The activity of F_1_-ATPase is determined by applying the Beer-Lambert law to calculate NADH oxidation in nmoles per minute per gram of protein. The assay was performed on chloroform-extracted F_1_-ATPase particles from the parental and from subunit α RNAi FL^WT^ and FL^PAS^ rescue cell lines induced with tetracycline for five days. Samples were incubated in an ATP hydrolysis buffer (50 mM Tris, pH 8.0, 50 mM KCl, 2 mM MgSO4, 200 µM NADH, 1 mM PEP, 2 mM MgATP, lactate dehydrogenase (5µl/ml, Sigma) and pyruvate kinase (5µl/ml, Sigma), and absorbance was measured for three minutes using a UV-visible spectrophotometer. Non-specific ATPase activity was subtracted from a 1 mM azide preincubation and F_1_-ATPase specific activity was calculated based on a standard curve [57].

### ATP production assay

A mitochondrial ATP production assay was performed on day 5 induced RNAi rescue cell lines as described [56]. The organellar fraction was isolated by extraction with 0.015% digitonin and resuspended in ATP production assay buffer. To assess maximum ATP production from F_1_F_o_-ATP synthase, control samples were preincubated with inhibitors of complex III (0.2 µM antimycin) and complex IV (1 mM KCN). Samples were incubated with two substrates, glycerol-3-phosphate (5 mM) and ADP (67 µM), and the reaction was terminated after 20 minutes with perchloric acid. After centrifugation and neutralization, ATP concentration was measured using the Roche ATP Bioluminescence Assay Kit on a Tecan Spark plate reader, and luminescence values were analyzed with GraphPad Prism.

### Immunofluorescence assay

A total of 2×10^7^ cells were harvested and incubated in SDM-79 medium with 200 nM Mitotracker Red CMXRos for 30 minutes. After fixation with 3.7% formaldehyde, cells were applied to coverslips, washed in PBS, permeabilized (0.1% Triton X-100/PBS), and blocked (5.5 % FBS/PBS-T). They were then incubated with primary mAb-V5 antibody (1:2000 dilution), followed by a secondary Alexa Fluor 488-conjugated antibody (1:400 dilution). Coverslips were mounted with ProLong Glass Antifade Mountant with NucBlue, and images were captured using an Axioplan 2 imaging fluorescent microscope equipped with an Olympus DP73 CCD camera.

### Subcellular fractionation using digitonin

Total of 1×10^8^ cells were harvested, washed and treated with 0.015% (w/v) digitonin on ice for 5 min. The cells were centrifuged at 7000 rpm for 3 min at 4°C, and the cytosolic supernatant was collected. The organellar pellet was resuspended in the same volume as the cytosolic volume of PBS. Both fractions were prepared for SDS-PAGE western blot analysis. To standardize the digitonin concentration for the release of cytosolic, glycosomal and mitochondrial proteins from *T. brucei*, a digitonin latency experiment was performed as described earlier [60]. For the FRET assay, three digitonin concentrations were chosen to isolate the cytosolic and mitochondrial fractions. Briefly, 6.5×10^8^ *T. brucei* cells were collected, rinsed twice with PBS, and resuspended in STEN buffer (250 mM sucrose, 25 mM Tris-HCl pH-7.4, 150 mM NaCl, 1 mM DTT) to 10 mg protein/ml. 1 mg of protein were incubated with 0.02 mg digitonin at 25°C for 4 min, then centrifuged at 14000xg for 2 min. The supernatant (cytosolic content) was collected and the pellet was treated with 0.16 mg digitonin/mg protein to extract glycosomal material. This procedure was repeated again to obtain mitochondrial material with 0.4 mg digitonin/mg protein. The protein concentration of the cytosolic and mitochondrial fractions was determined by BCA assay and used for FRET assay.

### FRET proteolytic assay with cytosolic and mitochondrial fractions

For the in vitro FRET assay, 50 ug of cytosolic and mitochondrial proteins was pre-treated with with 1 μM bestatin and 1 μM benzylsuccinic acid for 20 min at room temperature to inhibit some aminopeptidases and carboxypeptidase A, respectively. Subsequently, 32 μM of FRET peptides 1 and 2 were added (GenScript, USA), and the plate was immediately moved to a TECAN device. The fluorescence intensity was monitored at 490 nm over a period of 60 min.

---

**Supplemental Figure S1.**
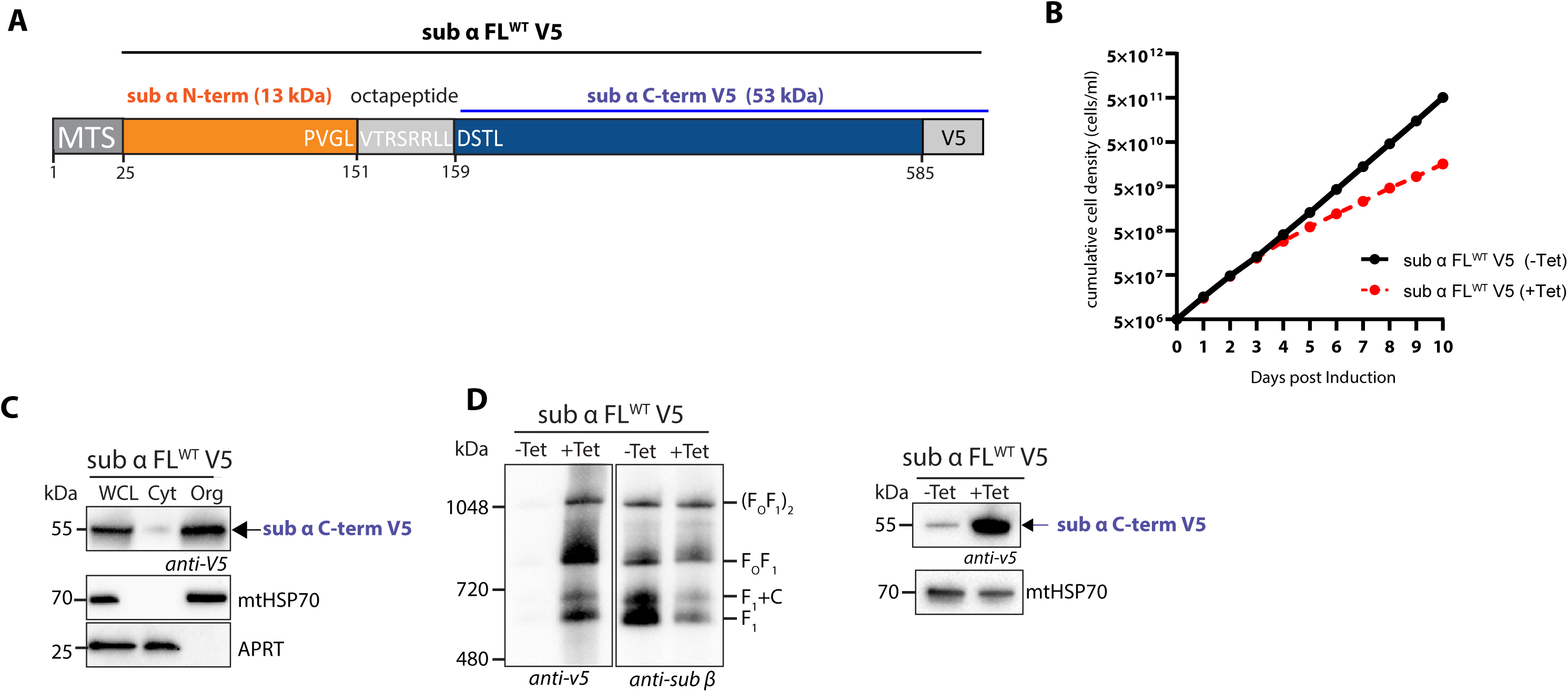
Inducible expression of subunit α FL^WT^ V5 causes a growth reduction in *T. brucei.* **A.** Schematic of *T. brucei* subunit α FL^WT^ with a C-terminal V5 tag. **B**. Growth curves of noninduced (-Tet) and induced (+Tet) cells expressing subunit α FL^WT^ V5. **C**. Western blot analysis of whole cell lysate (WCL), cytosol (Cyt) and digitonin-extracted enriched organellar fractions (Org) from *T. brucei* expressing subunit α FL^WT^ tagged with a V5 epitope. The relevant sizes of the protein marker are indicated on the left. Antibodies recognizing mitochondrial and cytosolic markers, mtHSP70 and APRT, respectively, were used as controls for proper fractionation. **D.** Representative immunoblots of mitochondrial lysates from *T. brucei* subunit α FL^WT^ V5 cell lines either noninduced (-Tet) or induced for 2 days (+Tet) were resolved by BN-PAGE and probed with antibodies against the V5 epitope and F_1_-ATPase subunit β. Positions of high molecular weight marker are shown. **E.** Mitochondrial lysates from **D** were separated on SDS PAGE and probed with antibodies against V5. Immunoblot with anti-mtHSP70 antibody served as a loading control.

**Supplemental Figure S2.**
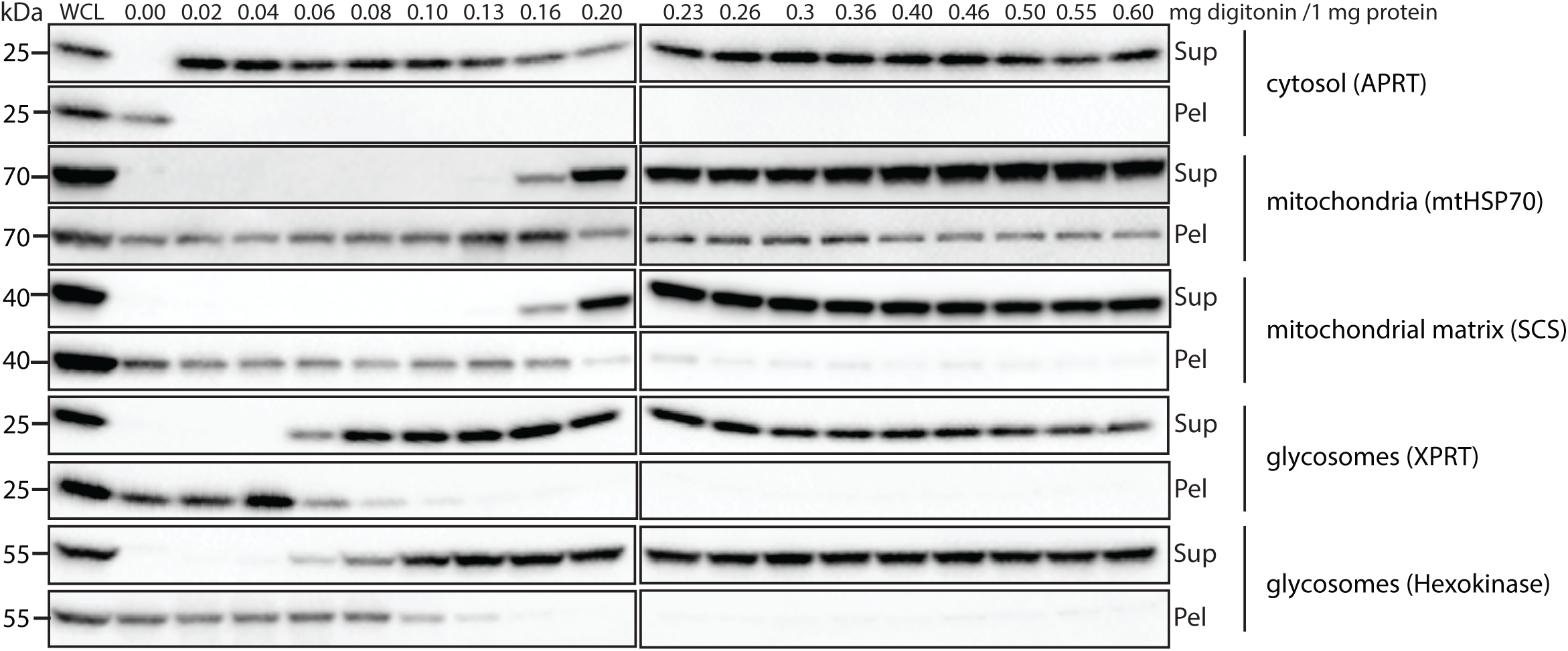
Digitonin latency of *T. brucei* PF427 cells. Immunoblots of whole cell lysates (WCL) and subcellular fractions obtained from treatments with digitonin at increasing concentrations. Antibodies recognizing specific proteins localized in either the cytosol, mitochondrion, mitochondrial matrix or glycosomes were applied. Protein marker sizes are indicated on the left.

**Supplemental Table S1. *T. brucei* transgenic cell lines generated in this study.**

**Supplemental Table S2. List of oligonucleotides used in this study.**

**Supplemental Table S3. List of used antibodies in this study.**

**Supplemental Table S4. Raw data supporting Figure 4E and 4F analysis.**

## AUTHOR CONTRIBUTIONS STATEMENT

M.J. performed the majority of the experimental work, analyzed data, and prepared figures. B.P. supervised the project, analyzed data, and wrote the manuscript. K.S. contributed experimentally at the beginning of the project. A.S. supervised the project, revised the manuscript, and provided funding. A.Z. supervised the project, revised the manuscript, and provided funding.

## ACKNOWLEDGEMENTS

We would like to thank Martina Slapničková for excellent technical support, Anzhelika Butenko for supplying us with the *Azumiobodo hoyamushi* and *Neobodo designis* α subunit protein sequences used in the euglenozoa alignments. We also appreciate Prashant Chauhan patiently providing us with a tutorial on how to use the UCSF ChimeraX software. We also acknowledge that the molecular graphics and analyses performed with UCSF ChimeraX were developed by the Resource for Biocomputing, Visualization, and Informatics at the University of California, San Francisco. Finally, we are grateful to Ken Stuart for the never-ending gift of the hybridoma cell line expressing the mtHSP70 monoclonal antibody.

## REFERENCES

1. Yanagisawa S, Frasch WD (2017) Protonation-dependent stepped rotation of the F-type ATP synthase c-ring observed by single-molecule measurements. J Biol Chem 292: 17093–17100.

2. Abrahams JP, Leslie AG, Lutter R, Walker JE (1994) Structure at 2.8 A resolution of F1-ATPase from bovine heart mitochondria. Nature 370: 621–628.

3. Boyer PD, Chance B, Ernster L, Mitchell P, Racker E, et al. (1977) Oxidative phosphorylation and photophosphorylation. Annu Rev Biochem 46: 955–966.

4. Wilkens S, Capaldi RA (1998) ATP synthase’s second stalk comes into focus. Nature 393: 29.

5. Watt IN, Montgomery MG, Runswick MJ, Leslie AG, Walker JE (2010) Bioenergetic cost of making an adenosine triphosphate molecule in animal mitochondria. Proc Natl Acad Sci U S A 107: 16823–16827.

6. Stock D, Leslie AG, Walker JE (1999) Molecular architecture of the rotary motor in ATP synthase. Science 286: 1700–1705.

7. Ferguson SJ (2010) ATP synthase: from sequence to ring size to the P/O ratio. Proc Natl Acad Sci U S A 107: 16755–16756.

8. Vazquez-Acevedo M, Vega-deLuna F, Sanchez-Vasquez L, Colina-Tenorio L, Remacle C, et al. (2016) Dissecting the peripheral stalk of the mitochondrial ATP synthase of chlorophycean algae. Biochim Biophys Acta 1857: 1183–1190.

9. Zikova A, Schnaufer A, Dalley RA, Panigrahi AK, Stuart KD (2009) The F(0)F(1)-ATP synthase complex contains novel subunits and is essential for procyclic Trypanosoma brucei. PLoS Pathog 5: e1000436.

10. Perez E, Lapaille M, Degand H, Cilibrasi L, Villavicencio-Queijeiro A, et al. (2014) The mitochondrial respiratory chain of the secondary green alga Euglena gracilis shares many additional subunits with parasitic Trypanosomatidae. Mitochondrion.

11. Yadav KNS, Miranda-Astudillo HV, Colina-Tenorio L, Bouillenne F, Degand H, et al. (2017) Atypical composition and structure of the mitochondrial dimeric ATP synthase from Euglena gracilis. Biochim Biophys Acta Bioenerg 1858: 267–275.

12. Kuhlbrandt W (2019) Structure and Mechanisms of F-Type ATP Synthases. Annu Rev Biochem 88: 515–549.

13. Muhleip AW, Joos F, Wigge C, Frangakis AS, Kuhlbrandt W, et al. (2016) Helical arrays of U-shaped ATP synthase dimers form tubular cristae in ciliate mitochondria. Proc Natl Acad Sci U S A 113: 8442–8447.

14. Strauss M, Hofhaus G, Schroder RR, Kuhlbrandt W (2008) Dimer ribbons of ATP synthase shape the inner mitochondrial membrane. EMBO J 27: 1154–1160.

15. Davies KM, Strauss M, Daum B, Kief JH, Osiewacz HD, et al. (2011) Macromolecular organization of ATP synthase and complex I in whole mitochondria. Proc Natl Acad Sci U S A 108: 14121–14126.

16. Davies KM, Daum B, Kuhlbrandt W, Anselmi C, Faraldo-Gomez J (2012) Structure of the mitochondrial ATP synthase and its role in shaping mitochondria cristae. Microsc Microanal 18 Suppl 2: 56–57.

17. Dudkina NV, Oostergetel GT, Lewejohann D, Braun HP, Boekema EJ (2010) Row-like organization of ATP synthase in intact mitochondria determined by cryo-electron tomography. Biochim Biophys Acta 1797: 272–277.

18. Blum TB, Hahn A, Meier T, Davies KM, Kuhlbrandt W (2019) Dimers of mitochondrial ATP synthase induce membrane curvature and self-assemble into rows. Proc Natl Acad Sci U S A.

19. Muhleip A, McComas SE, Amunts A (2019) Structure of a mitochondrial ATP synthase with bound native cardiolipin. Elife (Cambridge) 8.

20. Gahura O, Muhleip A, Hierro-Yap C, Panicucci B, Jain M, et al. (2022) An ancestral interaction module promotes oligomerization in divergent mitochondrial ATP synthases. Nat Commun 13: 5989.

21. Muhleip AW, Dewar CE, Schnaufer A, Kuhlbrandt W, Davies KM (2017) In situ structure of trypanosomal ATP synthase dimer reveals a unique arrangement of catalytic subunits. Proc Natl Acad Sci U S A 114: 992–997.

22. Davies KM, Anselmi C, Wittig I, Faraldo-Gomez JD, Kuhlbrandt W (2012) Structure of the yeast F1Fo-ATP synthase dimer and its role in shaping the mitochondrial cristae. Proc Natl Acad Sci U S A 109: 13602–13607.

23. Dudkina NV, Kouril R, Bultema JB, Boekema EJ (2010) Imaging of organelles by electron microscopy reveals protein-protein interactions in mitochondria and chloroplasts. FEBS Lett 584: 2510–2515.

24. Munoz-Gomez SA, Slamovits CH, Dacks JB, Baier KA, Spencer KD, et al. (2015) Ancient homology of the mitochondrial contact site and cristae organizing system points to an endosymbiotic origin of mitochondrial cristae. Curr Biol 25: 1489–1495.

25. Panek T, Elias M, Vancova M, Lukes J, Hashimi H (2020) Returning to the Fold for Lessons in Mitochondrial Crista Diversity and Evolution. Curr Biol 30: R575–R588.

26. Seravin LN (1993) [The basic types and forms of the fine structure of mitochondrial cristae: the degree of their evolutionary stability (capacity for morphological transformations)]. Tsitologiia 35: 3–34.

27. Montgomery MG, Gahura O, Leslie AGW, Zikova A, Walker JE (2018) ATP synthase from Trypanosoma brucei has an elaborated canonical F1-domain and conventional catalytic sites. Proc Natl Acad Sci U S A.

28. Davies KM, Kuhlbrandt W (2018) Structure of the catalytic F1 head of the F1-Fo ATP synthase from Trypanosoma brucei. Proc Natl Acad Sci U S A.

29. Alvarez-Jarreta J, Amos B, Aurrecoechea C, Bah S, Barba M, et al. (2024) VEuPathDB: the eukaryotic pathogen, vector and host bioinformatics resource center in 2023. Nucleic Acids Res 52: D808–D816.

30. Aslett M, Aurrecoechea C, Berriman M, Brestelli J, Brunk BP, et al. (2010) TriTrypDB: a functional genomic resource for the Trypanosomatidae. Nucleic Acids Res 38: D457–462.

31. Shanmugasundram A, Starns D, Bohme U, Amos B, Wilkinson PA, et al. (2023) TriTrypDB: An integrated functional genomics resource for kinetoplastida. PLoS Negl Trop Dis 17: e0011058.

32. Coustou V, Biran M, Breton M, Guegan F, Riviere L, et al. (2008) Glucose-induced remodeling of intermediary and energy metabolism in procyclic Trypanosoma brucei. J Biol Chem 283: 16342–16354.

33. Lamour N, Riviere L, Coustou V, Coombs GH, Barrett MP, et al. (2005) Proline metabolism in procyclic Trypanosoma brucei is down-regulated in the presence of glucose. J Biol Chem 280: 11902–11910.

34. Dewar CE, Casas-Sanchez A, Dieme C, Crouzols A, Haines LR, et al. (2022) Oxidative Phosphorylation Is Required for Powering Motility and Development of the Sleeping Sickness Parasite Trypanosoma brucei in the Tsetse Fly Vector. MBio: e0235721.

35. Njogu RM, Whittaker CJ, Hill GC (1980) Evidence for a branched electron transport chain in Trypanosoma brucei. Mol Biochem Parasitol 1: 13–29.

36. Schnaufer A, Clark-Walker GD, Steinberg AG, Stuart K (2005) The F1-ATP synthase complex in bloodstream stage trypanosomes has an unusual and essential function. EMBO J 24: 4029–4040.

37. Bily T, Sheikh S, Mallet A, Bastin P, Perez-Morga D, et al. (2021) Ultrastructural Changes of the Mitochondrion During the Life Cycle of Trypanosoma brucei. J Eukaryot Microbiol 68: e12846.

38. Aphasizheva I, Aphasizhev R (2020) Mitochondrial RNA quality control in trypanosomes. Wiley Interdiscip Rev RNA: e1638.

39. Speijer D, Breek CK, Muijsers AO, Hartog AF, Berden JA, et al. (1997) Characterization of the respiratory chain from cultured Crithidia fasciculata. Mol Biochem Parasitol 85: 171–186.

40. Maslov DA, Nawathean P, Scheel J (1999) Partial kinetoplast-mitochondrial gene organization and expression in the respiratory deficient plant trypanosomatid Phytomonas serpens. Mol Biochem Parasitol 99: 207–221.

41. Brown BS, Stanislawski A, Perry QL, Williams N (2001) Cloning and characterization of the subunits comprising the catalytic core of the Trypanosoma brucei mitochondrial ATP synthase. Mol Biochem Parasitol 113: 289–301.

42. Nelson RE, Aphasizheva I, Falick AM, Nebohacova M, Simpson L (2004) The I-complex in Leishmania tarentolae is an uniquely-structured F(1)-ATPase. Mol Biochem Parasitol 135: 221–224.

43. Gahura O, Subrtova K, Vachova H, Panicucci B, Fearnley IM, et al. (2018) The F1 -ATPase from Trypanosoma brucei is elaborated by three copies of an additional p18-subunit. FEBS J 285: 614–628.

44. Meng EC, Goddard TD, Pettersen EF, Couch GS, Pearson ZJ, et al. (2023) UCSF ChimeraX: Tools for structure building and analysis. Protein Sci 32: e4792.

45. Schechter I, Berger A (1967) On the size of the active site in proteases. I. Papain. Biochem Biophys Res Commun 27: 157–162.

46. Schechter I, Berger A (1968) On the active site of proteases. 3. Mapping the active site of papain; specific peptide inhibitors of papain. Biochem Biophys Res Commun 32: 898–902.

47. Spikes TE, Montgomery MG, Walker JE (2020) Structure of the dimeric ATP synthase from bovine mitochondria. Proc Natl Acad Sci U S A.

48. Srivastava AP, Luo M, Zhou W, Symersky J, Bai D, et al. (2018) High-resolution cryo-EM analysis of the yeast ATP synthase in a lipid membrane. Science 360.

49. Crooks GE, Hon G, Chandonia JM, Brenner SE (2004) WebLogo: a sequence logo generator. Genome Res 14: 1188–1190.

50. Jumper J, Evans R, Pritzel A, Green T, Figurnov M, et al. (2021) Highly accurate protein structure prediction with AlphaFold. Nature 596: 583–589.

51. Varadi M, Bertoni D, Magana P, Paramval U, Pidruchna I, et al. (2024) AlphaFold Protein Structure Database in 2024: providing structure coverage for over 214 million protein sequences. Nucleic Acids Res 52: D368–D375.

52. Stein V, Nabi M, Alexandrov K (2017) Ultrasensitive Scaffold-Dependent Protease Sensors with Large Dynamic Range. ACS Synth Biol 6: 1337–1342.

53. Schlapschy M, Binder U, Borger C, Theobald I, Wachinger K, et al. (2013) PASylation: a biological alternative to PEGylation for extending the plasma half-life of pharmaceutically active proteins. Protein Eng Des Sel 26: 489–501.

54. Meyer B, Wittig I, Trifilieff E, Karas M, Schagger H (2007) Identification of two proteins associated with mammalian ATP synthase. Mol Cell Proteomics 6: 1690–1699.

55. Subrtova K, Panicucci B, Zikova A (2015) ATPaseTb2, a unique membrane-bound FoF1-ATPase component, is essential in bloodstream and dyskinetoplastic trypanosomes. PLoS Pathog 11: e1004660.

56. Allemann N, Schneider A (2000) ATP production in isolated mitochondria of procyclic Trypanosoma brucei. Mol Biochem Parasitol 111: 87–94.

57. Gahura O, Panicucci B, Vachova H, Walker JE, Zikova A (2018) Inhibition of F1 -ATPase from Trypanosoma brucei by its regulatory protein inhibitor TbIF1. FEBS J.

58. Gahura O, Zikova A (2019) Isolation of F1-ATPase from the Parasitic Protist Trypanosoma brucei. J Vis Exp.

59. Pullman ME, Penefsky HS, Datta A, Racker E (1960) Partial resolution of the enzymes catalyzing oxidative phosphorylation. I. Purification and properties of soluble dinitrophenol-stimulated adenosine triphosphatase. J Biol Chem 235: 3322–3329.

60. Riviere L, Moreau P, Allmann S, Hahn M, Biran M, et al. (2009) Acetate produced in the mitochondrion is the essential precursor for lipid biosynthesis in procyclic trypanosomes. Proc Natl Acad Sci U S A 106: 12694–12699.

61. Moyersoen J, Choe J, Kumar A, Voncken FG, Hol WG, et al. (2003) Characterization of Trypanosoma brucei PEX14 and its role in the import of glycosomal matrix proteins. Eur J Biochem 270: 2059–2067.

62. Elustondo P, Martin LA, Karten B (2017) Mitochondrial cholesterol import. Biochim Biophys Acta Mol Cell Biol Lipids 1862: 90–101.

63. Ashworth LA, Green C (1966) Plasma membranes: phospholipid and sterol content. Science 151: 210–211.

64. Yeagle PL (1985) Cholesterol and the cell membrane. Biochim Biophys Acta 822: 267–287.

65. Coustou V, Guegan F, Plazolles N, Baltz T (2010) Complete in vitro life cycle of Trypanosoma congolense: development of genetic tools. PLoS Negl Trop Dis 4: e618.

66. Stuart KD (1971) Evidence for the retention of kinetoplast DNA in an acriflavine-induced dyskinetoplastic strain of Trypanosoma brucei which replicates the altered central element of the kinetoplast. J Cell Biol 49: 189–195.

67. Flaspohler JA, Jensen BC, Saveria T, Kifer CT, Parsons M (2010) A novel protein kinase localized to lipid droplets is required for droplet biogenesis in trypanosomes. Eukaryot Cell 9: 1702–1710.

68. Wirtz E, Leal S, Ochatt C, Cross GA (1999) A tightly regulated inducible expression system for conditional gene knock-outs and dominant-negative genetics in Trypanosoma brucei. Mol Biochem Parasitol 99: 89–101.

69. Trotter JR, Ernst NL, Carnes J, Panicucci B, Stuart K (2005) A deletion site editing endonuclease in Trypanosoma brucei. Mol Cell 20: 403–412.

70. Schneider A, Charriere F, Pusnik M, Horn EK (2007) Isolation of mitochondria from procyclic Trypanosoma brucei. Methods Mol Biol 372: 67–80.

71. Walker JE, Fearnley IM, Gay NJ, Gibson BW, Northrop FD, et al. (1985) Primary structure and subunit stoichiometry of F1-ATPase from bovine mitochondria. J Mol Biol 184: 677–701.

72. Hierro-Yap C, Subrtova K, Gahura O, Panicucci B, Dewar C, et al. (2021) Bioenergetic consequences of FoF1-ATP synthase/ATPase deficiency in two life cycle stages of Trypanosoma brucei. J Biol Chem: 100357.

73. Wittig I, Carrozzo R, Santorelli FM, Schagger H (2007) Functional assays in high-resolution clear native gels to quantify mitochondrial complexes in human biopsies and cell lines. Electrophoresis 28: 3811–3820.

74. Kostygov AY, Karnkowska A, Votypka J, Tashyreva D, Maciszewski K, et al. (2021) Euglenozoa: taxonomy, diversity and ecology, symbioses and viruses. Open Biol 11: 200407.

